# T2T Genome and Population Resequencing Reveal *OfCCD4* Alleles Orchestrate Petal Color and Scent in *Osmanthus fragrans*

**DOI:** 10.64898/2026.02.14.705464

**Authors:** Shengjun Liu, Shiling You, Jinmei Yuan, Xumei Zeng, Qinglian Yang, Shuyi Xu, Wan Xi, Zesi Peng, Linlin Zhu, Linlin Zhong, Ya-fei Tan, Ri-Ru Zheng

## Abstract

*Osmanthus fragrans* is prized for its floral color and fragrance, both key targets for genetic improvement. However, the lack of a complete genome assembly and comprehensive population structure hinders gene dissection and marker development for breeding. To investigate the genetic basis of color–scent interaction, a telomere-to-telomere (T2T) genome assembly is generated, and whole-genome resequencing of 100 cultivars is performed. Integrative population and metabolomic analyses reveal a trade-off: orange-red (‘Aurantiacus’) cultivars accumulate high α/β-carotene but low aromatic apocarotenoids (α/β-ionone), while yellow-white cultivars show the opposite pattern. Divergence mapping identifies *OfCCD4* as the major underlying locus. Three alleles are characterized—functional (A), partially functional (a^Del^), and a frameshift null (a^Stop^)—with a^Stop^ strictly co-segregating with the ‘Aurantiacus’ phenotype. Transient and stable transformations confirm that A and a^Del^ cleave α/β-carotene into α/β-ionone, while a^Stop^ abolishes this activity. A co-dominant PCR marker is developed based on allele-specific polymorphisms. *OfCCD4* is thus established as the key regulator of the color–scent trade-off in osmanthus. The identified alleles and molecular marker enable rapid, low-cost genotyping at the seedling stage—offering particular value for marker-assisted selection in ornamental breeding, where extended juvenility makes phenotypic selection highly time- and resource-intensive.

## 1. Introduction

*Osmanthus fragrans* is a world-renowned ornamental and aromatic plant, highly valued for its floral color and intense fragrance, with wide applications in food, cosmetics, and healthcare products^[0,0]^. Cultivars are traditionally classified into four groups based on petal color and flowering habit: the ‘Albus’ (light yellow-white), ‘Luteus’ (yellow), ‘Aurantiacus’ (orange-red), and ‘Asiaticus’ (pale yellow, multi-seasonal) groups^[0,0]^. Notably, there is a striking inverse correlation between petal color and scent production: orange-red ‘Aurantiacus’ cultivars accumulate high levels of carotenoid pigments (e.g., α-carotene and β-carotene) but produce low amounts of aromatic apocarotenoids (e.g., β-ionone and α-ionone), whereas yellow-white non-‘Aurantiacus’ cultivars exhibit the opposite pattern^[0-0]^. This natural variation presents an ideal system to dissect the genetic regulators governing the trade-off between pigment-based coloration and scent production.

Carotenoids and their enzymatic cleavage products, apocarotenoids, are central to this trade-off. Carotenoids serve as essential pigments, providing yellow to red hues ^[0]^, while their cleavage by carotenoid cleavage dioxygenases (CCDs) generates volatile apocarotenoids that are key determinants of floral scent^[0,0]^. Specifically, *CCD4* has been recurrently identified as a major genetic determinant of color variation across diverse species. Loss-of-function mutations in *CCD4* lead to carotenoid hyperaccumulation, resulting in yellow or orange coloration in flowers, fruits, and tubers, as observed in chrysanthemum, peach, and potato^[0-0]^. This underscores *CCD4’*s conserved role as a metabolic switch balancing pigment retention and aroma production.

In *O. fragrans*, *OfCCD4* is known to catalyze the cleavage of carotenoids into ionones, and its expression is inversely correlated with carotenoid accumulation^[0,0,0]^. Transcription factors such as OfWRKY3 and OfMYB1R114 further regulate its expression, highlighting its central position in the color–scent network^[0,0]^. However, the fundamental genetic basis for the divergence between ‘Aurantiacus’ and non-‘Aurantiacus’ cultivars remains unresolved. Conflicting mechanisms have been proposed, focusing on promoter variants such as a 4-bp or 183-bp deletion, but these studies lacked comprehensive functional validation across a broad germplasm and did not establish a unified genetic model^[0-0]^.

While previous genome sequencing studies have significantly advanced research on *O. fragrans*^[0-0]^, the absence of a complete, gap-free assembly has hindered the precise characterization of structural variants and their functional implications. Recent progress in sequencing technologies and assembly algorithms has now made it feasible to generate complete, telomere-to-telomere (T2T) genomes^[0-0].^ Such T2T genomes have become foundational resources for functional genomics and molecular breeding in several key horticultural species^[0-0]^, highlighting their potential to resolve longstanding biological questions.

To address this gap, we constructed a high-quality T2T genome for *O. fragrans*, providing a complete reference for accurate variant discovery and structural annotation. Leveraging this resource, we integrated population resequencing with multi-trait phenotyping across 100 cultivars, which identified *OfCCD4* as the major locus governing floral color and scent variation. Functional assays confirmed that a frameshift allele (a^Stop^) results in carotenoid accumulation in orange-flowered ‘Aurantiacus’ cultivars, whereas functional alleles (A and aᴰᵉˡ) promote carotenoid cleavage and apocarotenoid biosynthesis in yellow/white-flowered ‘non-Aurantiacus’ cultivars. Our study thus resolves the genetic mechanism underlying the color–scent trade-off in osmanthus and reinforces *CCD4* as an evolutionarily conserved regulator of these traits. Furthermore, we developed a co-dominant PCR marker for rapid genotyping, offering a practical tool to accelerate precision breeding in this slow-cycling perennial.

## 2. Results

### 2.1. Strong inverse correlation between carotenoid accumulation and apocarotenoid biosynthesis in ‘Aurantiacus’ and non-‘Aurantiacus’ cultivars

To quantitatively assess the phenotypic variation between ‘Aurantiacus’ (orange-red petals) and non-‘Aurantiacus’ (yellow-white petals) cultivars (Fig. 1A), we performed comprehensive colorimetric and metabolic profiling on flowers from a diverse panel of 100 *O. fragrans* cultivars. Color quantification using the CIE L*a*b* color space revealed a clear bimodal distribution corresponding to the two traditional classifications. ‘Aurantiacus’ cultivars exhibited significantly lower L* (lightness) and b* (yellowness) values, but substantially higher a* (redness) values compared to non-‘Aurantiacus’ cultivars (Table S4). Principal component analysis (PCA) based on the combined phenotypic matrix (L*a*b* values) clearly separated the 100 cultivars into two distinct clusters, which aligned perfectly with the ‘Aurantiacus’ and non-‘Aurantiacus’ classifications (Fig. S3).

**Figure 1.**
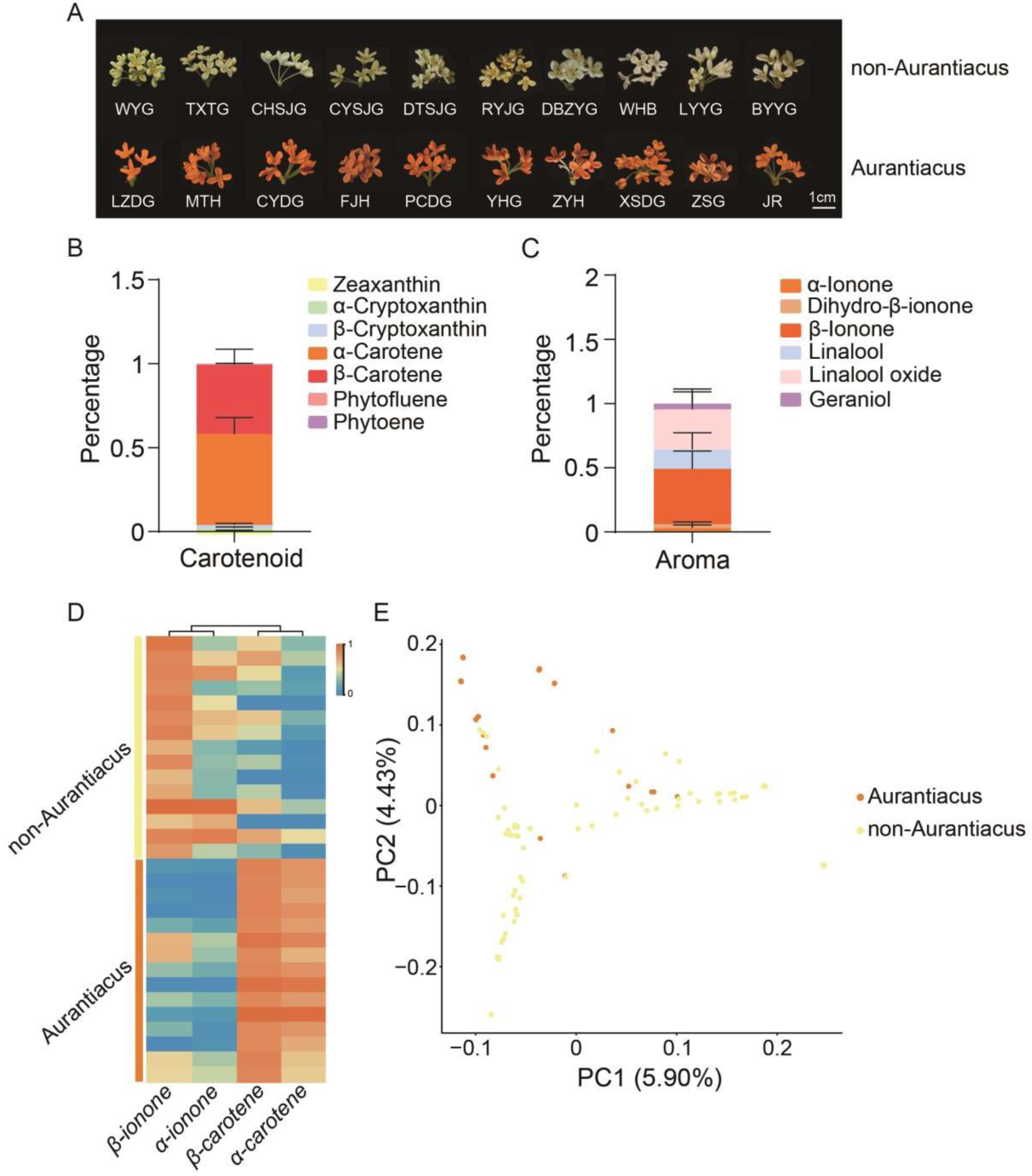
Carotenoid accumulation and aromatic apocarotenoid biosynthesis across 100 *Osmanthus fragrans* cultivars. (A) Representative flower phenotypes of typical ‘Aurantiacus’ (orange-flowered) and non-‘Aurantiacus’ (yellow/white-flowered) cultivars. Scale bar, 1 cm. (B) Average proportion of each carotenoid in petals of 15 ‘Aurantiacus’ cultivars. (C) Average proportion of each aromatic terpenoid in petals of 15 non-‘Aurantiacus’ cultivars. (D) Heatmap showing the relative content of α-carotene, β-carotene, α-ionone, and β-ionone in ‘Aurantiacus’ (n=15) versus non-‘Aurantiacus’ (n=15) groups. Data are normalized from 0 to 1. e, Principal component analysis of 24 ‘Aurantiacus’ cultivars, 76 non-‘Aurantiacus’ cultivars. Orange circles represent ‘Aurantiacus’; yellowish circles represent non-‘Aurantiacus’.

While osmanthus petals contain multiple pigment classes including carotenoids and flavonoids, prior metabolic profiling has established that the vivid orange-red hue of ‘Aurantiacus’ cultivars is predominantly due to elevated levels of carotenoids^[0]^. Due to the pronounced phenotypic differences between the two groups, we selected 30 representative cultivars for detailed carotenoid and apocarotenoid analysis (Table S5). In the current study, targeted metabolomic analysis identified seven carotenoids in *O. fragrans* petals: zeaxanthin, β-cryptoxanthin, α-cryptoxanthin, phytofluene, phytoene, α-carotene, and β-carotene, with α-carotene and β-carotene being the dominant compounds reach to 95% (Fig. 1B). More details, the average proportions of α-carotene and β-carotene were 21.8- and 19.3-fold higher, respectively, in ‘Aurantiacus’ group compared to non-‘Aurantiacus’ group(Fig. 1B; Fig. S4A). Concurrent profiling of aroma compounds via GC-MS (Gas Chromatography-Mass Spectrometry) identified the combined abundance of α-ionone, β-ionone, and dihydro-β-ionone represented the predominant fraction of aroma-active terpenoids reach to 46%, particularly in non-‘Aurantiacus’ cultivars (Fig. 1C; Fig. S4B). Furthermore, the concentrations of all three apocarotenoids were significantly higher in non-‘Aurantiacus’ cultivars, with average levels exceeding those in ‘Aurantiacus’ cultivars by 6.14 fold (Fig. 1C; Fig. S4B; Table S5). This inverse relationship between pigment accumulation and aromatic apocarotenoid content reveals a metabolic trade-off at the population level (Fig. 1D).

In order to identify the the major locus for controlling color and fragrance in *Osmanthus fragrans*, we collected and sequenced 100 cultivars (Table S6). We performed population genomic analysis using genome-wide SNP data obtained from resequencing the 100 cultivars. Neither Principal Component Analysis (PCA) of the SNP data revealed any significant population stratification corresponding to the ‘Aurantiacus’ and non-‘Aurantiacus’ groups (Fig. 1E). This indicates that the dramatic difference in color and scent is not a consequence of broad genome-wide genetic divergence or separate evolutionary lineages, but is likely controlled by a limited number of key genetic loci.

### 2.2. Chromosome-level T2T genome assembly of *O. fragrans ‘*Boye Yingui’

To further elucidate the genetic basis of petal color and scent in *Osmanthus fragrans*, we performed genome assembly for representative non-‘Aurantiacus’ (*O. fragrans ‘*Boye Yingui’). Combined PacBio high-fidelity (HiFi) reads and high-throughput/resolution chromosome conformation capture (Hi-C) data, we assembled a chromosome-level genome of *Osmanthus fragrans*. Then, we generated 39.33 Gb of Nanopore ultra-long reads to fill all the gaps for the chromosome-level genome assembly (Table S1). To further refine the assembly, MUMmer [0] and the previously published genome [0] were employed to order the 23 chromosomes (Fig. S1). The completeness of our genome assembly, as calculated by its Benchmarking Universal Single-Copy Orthologs (BUSCO) score, reached 98.9% (Fig. S2). Moreover, our assembly achieved the complete assembly of almost all chromosomal telomeres, except for the end of chromosome 6 and 19 (Fig. 2; Table S2). Notably, the newly assembled genome exhibited no gaps across all chromosomes, achieving a contig N_50_ of 29.90 Mb and the previously published genome with the contig N_50_ of 18.83%. Also, we annotated 54.88% transposable element (TE) and 37,359 genes in the newly assembled genome (Table 1). Among these, the LTR-Gypsy family accounts for the largest proportion, reaching 13.58% (Table S3).

**Figure 2.**
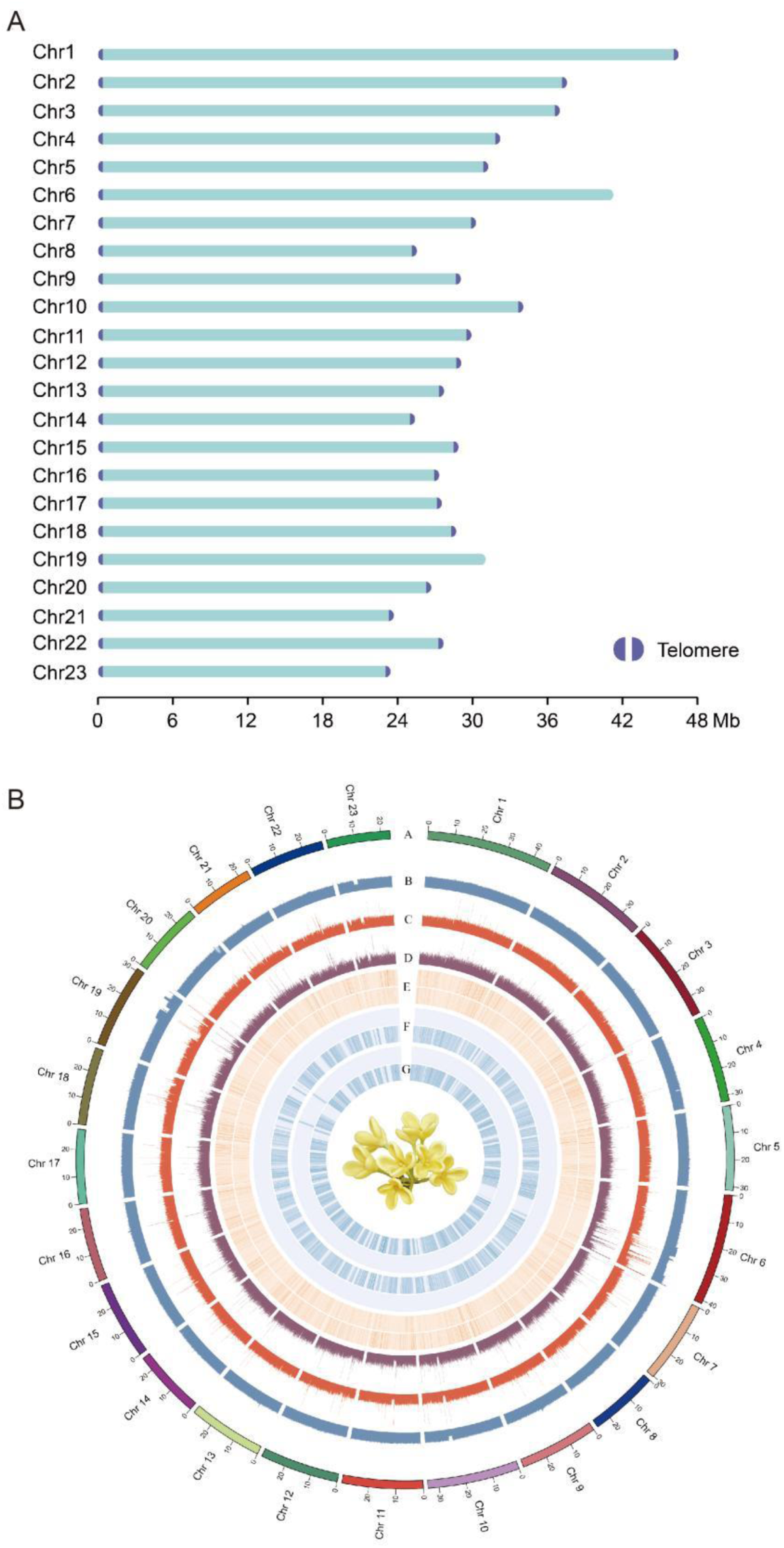
Telomere-to-telomere (T2T) genome assembly of *Osmanthus fragrans* ‘Boye Yingui’. (A) Purple semicircle, telomeres. (B) A circle: chromosome ideograms for T2T genome (Mb scale); B circle: GC content (%); C circle: Depth distribution of PacBio High-Fidelity reads; D circle: Depth distribution of nanopore Ultra-long reads; E circle: Complete comparison of BUSCO gene distribution in the genome, with the outer circle representing single-copy BUSCOs and the inner circle representing duplicated BUSCOs; F circle: The outer ring is the density distribution of homozygous SNPs, and the inner ring is the density distribution of heterozygous SNPs. G circle: The outer circle shows the distribution of homozygous InDel density, while the inner circle shows the distribution of heterozygous InDel density.

**Table 1.** T2T genome statistics for *Osmanthus fragrans*.

| Feature | *Osmanthus fragrans* |
| --- | --- |
| Assembled Size (Mb) | 701.58 |
| BUSCO completeness of assembly (%) | 98.9 |
| Number of gaps | 0 |
| Number of contigs | 23 |
| Largest contig (Mb) | 46.48 |
| Contig N<sub>50</sub> (Mb) | 29.9 |
| Percentage of transposable element (%) | 54.88 |
| Number of gene models | 37,359 |

### 2.3. Population genomic analysis identifies *OfCCD4* as the major locus governing color-fragrance trade-off

To investigate the population divergence between the ‘Aurantiacus’ population and non-‘Aurantiacus’ population, we computed pairwise genetic differentiation (*Fst*) values between 100 cultivars. We found that 11.35 Mb of the *Osmanthus fragrans* genomic regions were highly divergent relative to ‘Aurantiacus’ population and non-‘Aurantiacus’ population, a region harboring 537 genes (Table S7). The analysis pinpointed a highly significant locus associated with both petal color (α-carotene and β-carotene) and key aromatic apocarotenoid contents (α-ionone, β-ionone, dihydro-β-ionone). This locus colocalized precisely with the genomic region encoding *OfCCD4*, a known carotenoid cleavage dioxygenase, establishing it as the candidate responsible for the phenotypic divergence (Fig. 3A). Tissue-specific expression profiling further supported the central role of *OfCCD4*. The gene was predominantly expressed in floral tissues, with the highest transcript levels detected in petals (Fig. 3B; Table S8).

**Figure 3.**
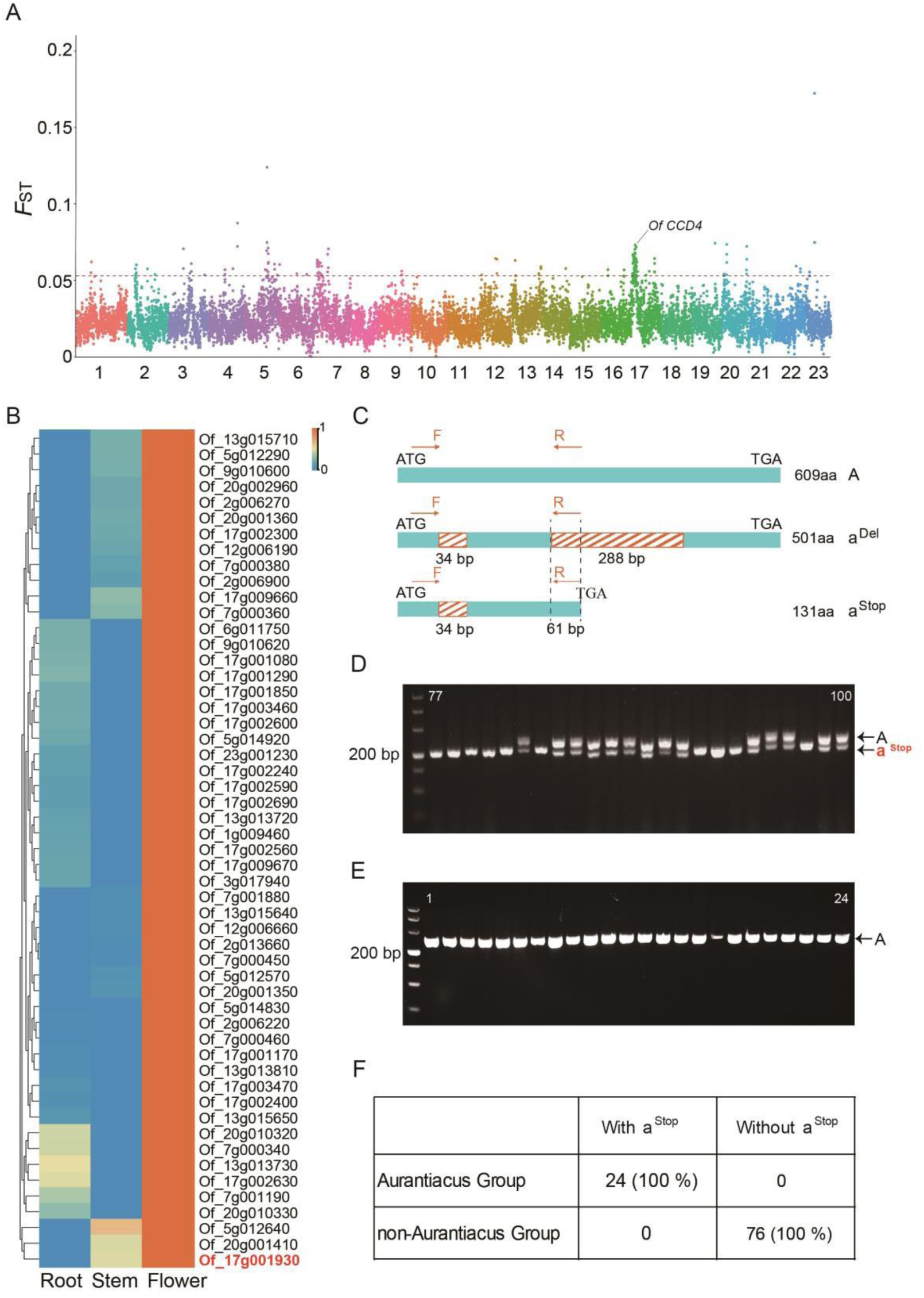
Population genomic analysis identifies *OfCCD4* as the major locus governing the color–fragrance trade-off. (A) Manhattan plot of genetic differentiation (*Fst*) across the genome between ‘Aurantiacus’ and non-‘Aurantiacus’ populations. Regions with *Fst* values in the top 1% (red dashed line) were considered significant selective sweeps. (B) Expression profiles of candidate genes from the significant region in different floral tissues; only flower-highly expressed candidates are shown. The star marks *OfCCD4*. (C) Schematic model of the three *OfCCD4* alleles: functional wild-type (A), a deletion variant (a^Del^), and a premature Stop codon variant (a^Stop^). Primer positions (F, forward; R, reverse) for allele-specific PCR are indicated. (D) Gel electrophoresis of PCR products for detecting the a^Stop^ and A/a^Del^ alleles in ‘Aurantiacus’ cultivars. Short band: a^Stop^; long band: A allele. (E) Representative PCR detection showing the presence of the A allele in non-‘Aurantiacus’ cultivars. (F) Summary of the distribution of *OfCCD4* coding sequence (cds) genotypes across 24 ‘Aurantiacus’ cultivars and 76 non-’Aurantiacus’ cultivars.

Given previous reports implicating *OfCCD4* promoter variants in its regulation^[0,0]^, we first analyzed the promoter sequences across our 100-cultivar panel. A previously reported 4-bp deletion near a CAAT-box did not segregate with flower color phenotype, as it was present in both ‘Aurantiacus’ and non-‘Aurantiacus’ groups (Fig. S5). We also developed a pair of primers to distinguish a widespread 183-bp deletion variant (designated P^Del^) from the normal promoter (P) (Fig. S6). Notably, the P^Del^ allele lacks several putative cis-regulatory elements, including hormone-responsive motifs (e.g., for MeJA, SA, and ABA), light-responsive elements (GA-motif, G-box), a WRKY-binding W-box, and a stress-related ARE motif (Table S9). This deletion was found in all genotype combinations (PP, PP^Del^, and P^Del^P^Del^) and was not specific to a color group, being present in both ‘Aurantiacus’ and non-‘Aurantiacus’ cultivars (Table S10). However, the absence of phenotypic specificity for these promoter variants across the population underscores that coding sequence variation, rather than promoter architecture, is the primary determinant of the functional output governing the color-scent trade-off (Table S10).

We next focused on the coding sequence of *OfCCD4* and identified three major allelic variants encoding structurally distinct proteins: (i) The allele *OfCCD4(A)* encodes a full-length protein of 609 amino acids (1,830 bp). (ii) The allele *OfCCD4^Stop^* (a^Stop^) carries 34 bp deletions, resulting in a frameshift that introduces a premature Stop codon; the predicted protein is truncated after 131 amino acids. (iii) The allele *OfCCD4^Del^*(a^Del^) contains 34 bp, and a 288 bp deletion. *OfCCD4a^Del^* exhibits a 287bp deletion compared to *OfCCD4a^Stop^*, with this deletion located at the translation termination site of *OfCCD4a^Stop^*, thereby allowing translation to continue and ultimately encoding 501 amino acids (Fig. 3c).

### 2.4. A co-dominant PCR marker system distinguishes ‘Aurantiacus’ and non-‘Aurantiacus’ cultivars

Based on population-wide haplotype analysis, we observed that the loss-of-function a^Stop^ allele was present in all 24 ‘Aurantiacus’ cultivars and absent in all 76 non-‘Aurantiacus’ cultivars. This perfect genotype–phenotype correlation confirms that coding sequence variation in *OfCCD4*, particularly the a^Stop^ allele, constitutes the key genetic determinant of the floral color–scent trade-off in *O. fragrans* (Fig. 3d-f; Fig. S7). In contrast, A and a^Del^ alleles were found in both groups but were not exclusively associated with either phenotype, indicating that they do not drive the major color–scent divergence (Table S11). Notably, all non-‘Aurantiacus’ cultivars exhibited a long amplicon in our co-dominant marker assay, corresponding to genotypes AA or A/a^Del^ (Fig. 3d-f), further supporting the diagnostic role of the a^Stop^ allele in distinguishing ‘Aurantiacus’ from non-‘Aurantiacus’ accessions.

To enable rapid and reliable genotyping, we developed a co-dominant PCR marker system that distinguishes the three *OfCCD4* alleles based on amplicon length polymorphism (Fig. 3c; Table S12). Population-wide screening using this marker revealed that all orange-red flowered ‘Aurantiacus’ cultivars exclusively carried the a^Stop^ allele with a short amplicon, while all yellow-white flowered non-‘Aurantiacus’ cultivars carried either the A allele (long amplicon) or the a^Del^ allele (no amplicon) (Fig. 3d-e; Fig. S7). This result perfectly aligns with the known coding sequence structures of each allele, underscoring the diagnostic power of the a^Stop^ allele for predicting the ‘Aurantiacus’ phenotype (Table S11).

### 2.5. Coding sequence variation in *OfCCD4* is the key determinant of enzymatic function and phenotype

To functionally validate the causal role of the three *OfCCD4* coding alleles, we conducted a series of *in planta* assays using both transient expression in the native *O. fragrans* petals and stable transformation in heterologous citrus callus systems (Fig. 4a; Table S12).

**Figure 4.**
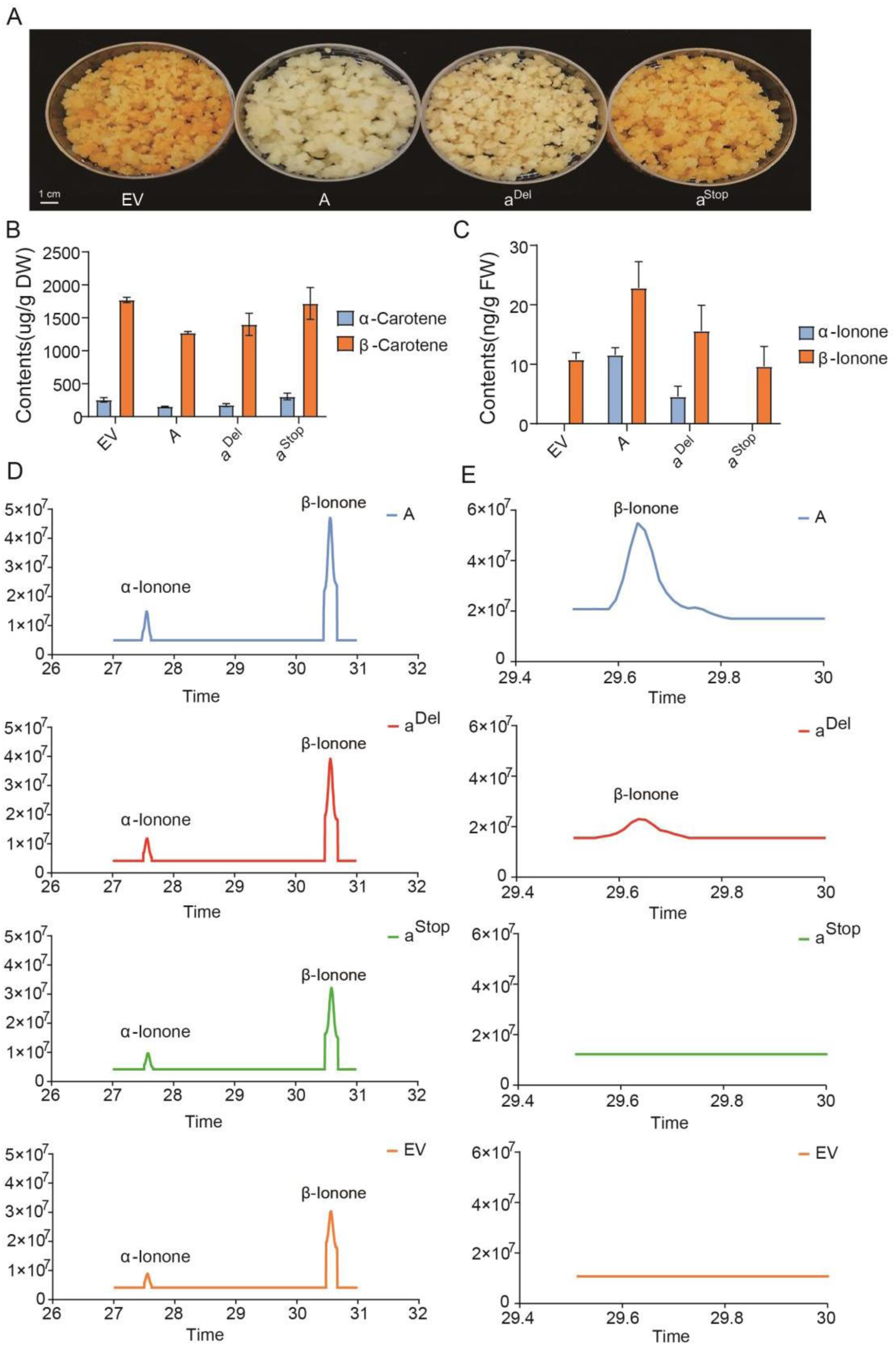
Functional verification of three *OfCCD4* alleles. (A) Stable overexpression assay in carotenoid-rich citrus callus overexpressing empty vector (EV) or different *OfCCD4* alleles. (B-C) Comparison of α-carotene, β-carotene, α-ionone, and β-ionone levels in *O. fragrans* petals transiently expressing EV and three different *OfCCD4* alleles. (D) Levels of α-ionone and β-ionone in the transgenic *O. fragrans* petals. (E) Levels of α-ionone and β-ionone in the transgenic citrus callus. Carotenoids were analyzed by HPLC, and aroma compounds were analyzed by GC-MS. Data represent mean ± SD of three biological replicates.

Transient overexpression of the functional A and partially functional a^Del^ alleles in petals of the carotenoid-rich ‘Aurantiacus’ cultivar ‘Gecheng Dangui’ resulted in a significant decrease in endogenous carotenoid content. Specifically, levels of α-carotene and β-carotene were reduced by an average of ∼40% and ∼28% in the A allele overexpression flowers compared to the empty vector control, respectively (Fig. 4b; Fig. S8). Concurrently, emission of the key apocarotenoid β-ionone increased by more than 2-fold (Fig. 4c-d; Fig. S9). In contrast, expression of the frameshift a^Stop^ allele produced no significant changes in either carotenoid or apocarotenoid levels, confirming its loss-of-function nature.

These findings were robustly corroborated through stable genetic transformation of citrus callus, a heterologous system with high carotenoid-accumulating capacity. Calli expressing the A or a^Del^ alleles displayed a marked reduction in total carotenoid content (predominantly β-carotene) and a corresponding increase in β-ionone production (Fig. 4e; Fig. S10-12). Conversely, lines expressing the a^Stop^ allele showed carotenoid profiles indistinguishable from the empty vector control and produced negligible apocarotenoids. Consistent with these metabolic shifts, callus infiltrated with the A or a^Del^ alleles exhibited visible reduction in orange pigmentation (Fig. 4a).

Collectively, these functional assays demonstrated a clear enzymatic hierarchy: the A allele possessed full cleavage activity; the a^Del^ allele retained partial but significant function; and the a^Stop^ allele was enzymatically null. This functional gradient directly translated to the observed phenotypes: alleles enabling carotenoid cleavage (A, a^Del^) led to pigment depletion and fragrance production (yellow-white, high-scent phenotype), while the non-functional a^Stop^ allele resulted in carotenoid retention and low fragrance (orange-red, low-scent phenotype) (Fig. 5). Therefore, we conclusively establish that coding sequence variation in *OfCCD4* is the primary genetic determinant governing the metabolic trade-off between petal color and floral fragrance in sweet osmanthus.

**Figure 5.**
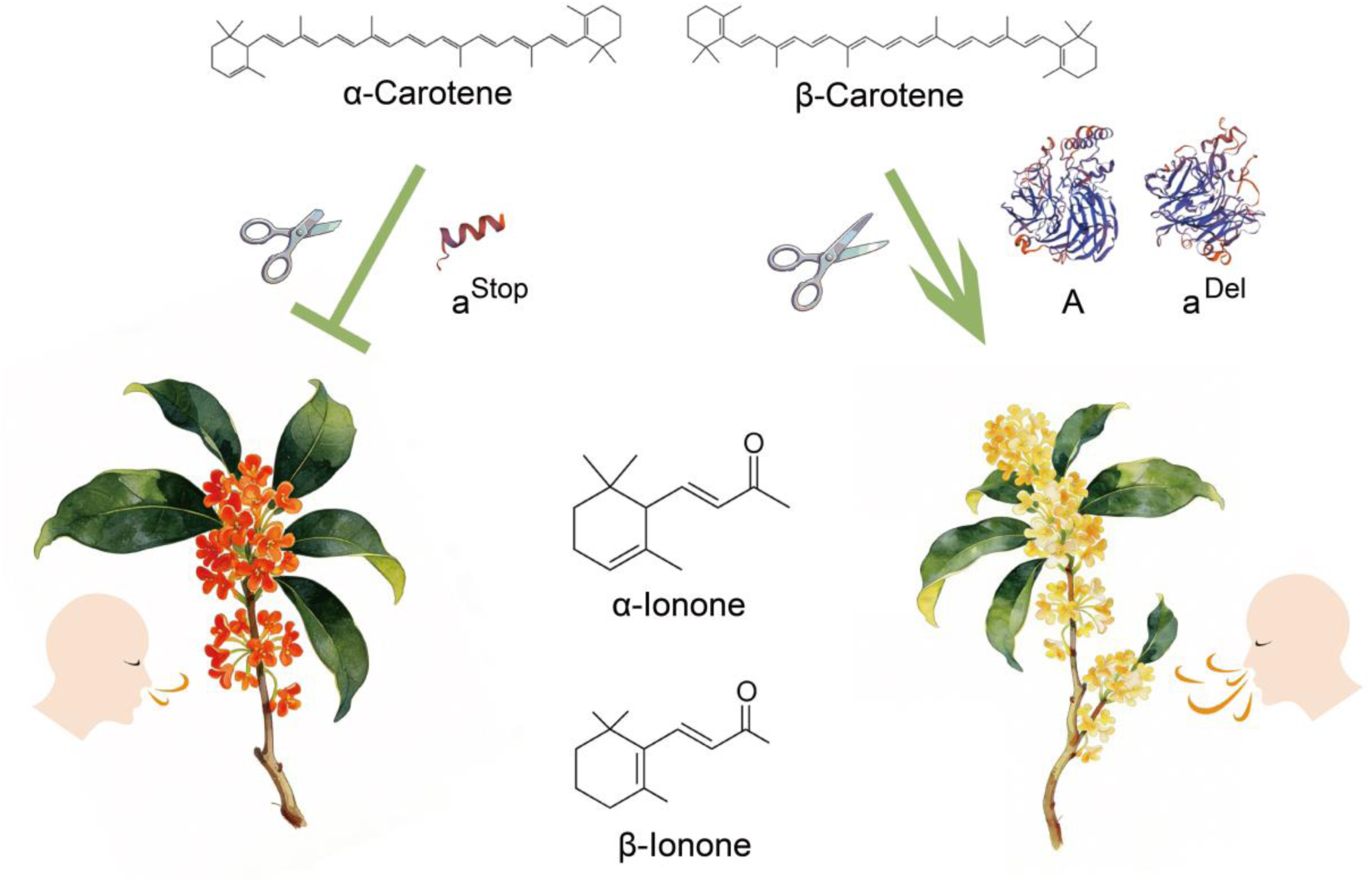
Model of *OfCCD4* as a genetic switch for manipulating color and fragrance traits. Schematic diagram illustrating the proposed mechanism. Functional *OfCCD4* (A allele) cleaves carotenoids (e.g., α-/β-carotene) to produce volatile apocarotenoids (e.g., α-/β-ionone), leading to fragrant but low-pigment (non-‘Aurantiacus’) flowers. Loss-of-function alleles (a^Del^, a^Stop^) impair this cleavage, allowing carotenoids to accumulate, resulting in deeply colored but low-fragrance (‘Aurantiacus’) flowers.

## 3. Discussion

Our study establishes *OfCCD4* as the central genetic determinant governing the classic trade-off between petal color and floral fragrance in *O. fragrans*. By integrating a high-quality T2T genome, population genomics, and functional validation, we provide a comprehensive genetic model that clarifies and extends previous findings, offering both mechanistic insights and practical tools for molecular-assisted breeding.

### 3.1. OfCCD4 as the central genetic switch governing the color–scent trade-off

The metabolic trade-off between color and scent in osmanthus is underpinned by distinct carotenoid and apocarotenoid profiles. Floral aroma is dominated by apocarotenoids such as β-ionone, dihydro-β-ionone, and α-ionone, which are key aroma-active compounds due to their high odor activity values and floral notes ^[0,0,0,0]^. In contrast, the orange-red coloration of ‘Aurantiacus’ cultivars is attributed to carotenoid accumulation, notably α-carotene and β-carotene^[0,0,0,0]^. Our results demonstrate that OfCCD4 acts as the enzymatic switch partitioning metabolic flux between these two compound classes, directly linking pigment accumulation with fragrance emission.

### 3.2. Coding Sequence Variation, Not Promoter Polymorphisms, Drives Phenotypic Divergence

*OfCCD4* coding sequence variation, rather than promoter polymorphisms, is the primary driver of phenotypic divergence in *O. fragrans.* Previous studies proposed conflicting mechanisms for flower color divergence, focusing on structural variants in the *OfCCD4* promoter ^[0,0]^. However, our population-wide genotyping revealed that neither the 183-bp deletion (P^Del^) nor the 4-bp CAAT-box deletion is strictly associated with the ‘Aurantiacus’ phenotype (Fig. S5, S6). Instead, we identified three coding alleles—functional (A), partially functional (a^Del^), and a frameshift null allele (a^Stop^)—with the a^Stop^ allele perfectly co-segregating with orange-red petals and low scent output (Fig. 3f). This finding underscores that coding sequence alterations, not regulatory variants, constitute the major genetic determinant of the color–scent trade-off in osmanthus.

### 3.3. Evolutionary Conservation of CCD4-Mediated Metabolic Trade-Offs Across Plant Families

The role of CCD4 in mediating color–scent trade-offs is evolutionarily conserved across diverse plant families^[0-0]^. For example, in *Brassica rapa*, a 15-bp deletion in *BrCCD4* causes a frameshift and yellow petals^[0]^; in *Brassica oleracea*, coding variants in *BoCCD4* determine petal color^[0]^. Loss-of-function alleles also lead to carotenoid accumulation in chrysanthemum^[0]^, potato^[0]^, and soybean^[0]^. These findings collectively show that coding sequence variants—frameshifts, premature stops, and missense mutations—directly impair CCD4 activity, promoting carotenoid retention and often reducing apocarotenoid emission. This conserved mechanism highlights CCD4’s role in balancing pigment and aroma across species.

### 3.4. Ecological and Adaptive Significance of ‘Aurantiacus’ Cultivars Under Northern Cultivation

The emergence of ‘Aurantiacus’ cultivars likely reflects adaptive advantages under northern cultivation. Historical records show that early varieties were light-colored before the Tang Dynasty (618–907 CE), with orange-red types appearing in the late Song Dynasty (960–1279 CE)^[0,0]^. This suggests human-mediated northward dispersal. Carotenoid accumulation from loss-of-function *OfCCD4* alleles may enhance cold tolerance by stabilizing membranes and reducing oxidative stress ^[0]^. Additionally, *OfCCD4* promoter contains multiple stress-responsive elements as other plants, implying expression modulation under stress ^[^**^错误!未找到引用源。^**^]^. The vivid orange color may also improve pollinator attraction in new environments ^[0]^. Thus, ‘Aurantiacus’ spread may combine molecular adaptation, physiological benefits, and reproductive success, illustrating how *CCD4* variation shapes both ornamental and ecological traits.

### 3.5. A Co-Dominant PCR Marker for Accelerated Molecular Breeding

A co-dominant PCR marker enables efficient selection for flower color and scent traits in breeding programs. Building on the diagnostic power of the a^Stop^ allele, we developed a robust molecular marker that reliably distinguishes the three *OfCCD4* alleles. This tool allows for early, non-destructive screening of seedlings, significantly shortening the breeding cycle for this slow-maturing perennial. Similar CCD4-based markers have been successfully deployed in strawberry, citrus, and peach for traits related to color, nutrition, and aroma ^[0-0]^, demonstrating the broad applicability of this approach in horticultural crop improvement.

In summary, our work establishes *OfCCD4* as the central genetic switch controlling the trade-off between carotenoid-based color and apocarotenoid-based fragrance in sweet osmanthus. By integrating T2T genomics, population genetics, and functional assays, we provide a unified model that reconciles prior discrepancies, reveals ecological correlates, and delivers a practical breeding tool. This study not only advances our understanding of color–scent coevolution in flowers but also demonstrates how genomic resources can bridge fundamental biology and applied horticulture, offering a replicable framework for the improvement of other aromatic and ornamental plants.

## 4. Experimental Section

### 4.1. Plant materials

A diverse panel of 100 *O. fragrans* cultivars, representing the three major horticultural groups (‘Aurantiacus’, ‘Albus’, ‘Luteus’ and ‘Asiaticus’). All germplasm materials were uniformly cultivated under standardized horticultural management in the nursery of Wuhan Xinzhou District Huaguoshan Ecological Park Agriculture Co., Ltd. (Wuhan, Hubei, China). Fully opened flowers from three biological replicates per cultivar were harvested for phenotypic and molecular analyses.

### 4.2. Telomere-to-telomere genome assembly and annotation

High-molecular-weight DNA was extracted from young leaves of the cultivar ‘Boye Yingui’ (‘Albus’ group) using the Nanobind Plant Nuclei Big DNA kit (Circulomics). Sequencing was performed on a PacBio Revio instrument (≥100× coverage) and an Illumina NovaSeq 6000 platform (≥50× coverage). Hi-C libraries were prepared and sequenced to scaffold contigs. HiFi reads were assembled using Hifiasm (v0.16.1)^[0]^ with default parameters. Nanopore ultralong reads were assembled using Nextdenovo (v2.5.2) (https://github.com/Nextomics/NextDenovo). The genome assembled by Nanopore reads was used to fill gaps of the genome assembled by Hifiasm using TGS-Gapcloser^[0]^. We also combined Hi-C reads to anchor contigs using 3d-dna pipeline (v180922). The plant telomeric sequences (CCCTAAA) were used to identify telomere in the assembled genome. Annotation was performed using a combined strategy: repetitive sequences were identified with Repeatmodeler2^[0]^ and RepeatMasker^[0]^; protein-coding genes were predicted via BRAKER3^[0-0]^, integrating RNA-seq data from five tissues and homology evidence from five related species.

### 4.3. Transcriptome sequencing of different tissues of *O. fragrans*

Root, stem and flower of *Osmanthus fragrans* ‘BYYG’ were collected for RNA-seq analysis. Three replicates of plant tissue were analyzed. For each sample, an RNA-seq library was constructed and sequenced on the Illumina platform (Table S8). All clean sequencing reads were aligned to the *Osmanthus fragrans* ‘BYYG’ T2T genome using Hisat2 (v2.2.1)^[0]^. The FPKM was quantified by Cufflinks (v2.2.1)^[0]^. Genes showing significantly different expression in flowers compared to root and stem were identified using the FPKM > 1, thresholds |log₂(fold change)| > 2 and p-value < 0.05.

### 4.4. Population resequencing and SNP calling

Whole-genome resequencing of the 100 cultivars was conducted on the Illumina NovaSeq 6000 platform (150 bp paired-end, ∼30×). Reads were aligned to the T2T reference genome using BWA (v0.7.17)^[0]^, duplicated mapping reads were removed using MarkDuplicates command from the GATK (v4.1.1)^[0]^. SNPs were called using HaplotypeCaller command from the GATK (v4.1.1). Raw VCF file were filtered using the Vcftools (v0.1.13) (--minDP 5 --max-missing 0.9 --maf 0.05)^[0]^. To analyse the genetic relationships among all 100 cultivars, we used all the filterd SNPs called through the above process performed PCA with default settings using GCTA (v1.26.0)^[0]^. We used PLINK (v1.90)^[0]^ for format conversion and using the R package *prcomp* to calculate PC1 and PC2.

Population differentiation was evaluated using *Fst*. Pairwise *Fst* values of comparisons between ‘Aurantiacus’ and non-‘Aurantiacus’ was determined using VCFtools with a 100-kb window and a 50-kb sliding window. Windows which containing the top 1% of *Fst* values in either comparisons of ‘Aurantiacus’ and non-‘Aurantiacus’ were identified as candidates for highly divergent regions.

### 4.5. Allele cloning, sequence analysis, and co-dominant marker development

Full-length cDNA of *OfCCD4* was amplified from petal total RNA using allele-specific primers, cloned into the pENTR™/D-TOPO® vector (Invitrogen), and Sanger-sequenced from at least three independent clones from typical cultivars (‘Aurantiacus’ cultivars: Zhuangyuanhong, Hongyan Ningxiang, Gecheng Dangui; non-‘Aurantiacus’ cultivars: Boye Yingui, Huangchuan Jingui, Fodingzhu). Sequence alignments were performed using Clustal Omega and visualized with GeneDoc. To distinguish the three major alleles (A, a^Stop^, and a^Del^) across the cultivar panel, we developed a co-dominant PCR marker system: the forward primer was designed upstream of the 34-bp deletion, and the reverse primer was placed within the 61-bp deletion region. PCR products were resolved on 3% agarose gels, yielding distinct banding patterns (A: long band; a^Stop^: short band; a^Del^: no product). Genotyping results were validated in a subset of cultivars by targeted amplicon sequencing.

### 4.6. Functional validation of *OfCCD4* alleles

The coding sequences of *OfCCD4* alleles (A, a^Stop^, a^Del^) were cloned into the plant expression vector pH7WG2D via Gateway recombination and transformed into *Agrobacterium tumefaciens EHA105*.

For transient expression in *O. fragrans* petals, *Agrobacterium* suspensions carrying the *OfCCD4a* allele constructs were infiltrated into petals of the carotenoid-rich cultivar ‘Gecheng Dangui’. After vacuum infiltration, petals were dark-incubated in 5% sucrose solution for 60 hours. Carotenoid extraction and quantification were performed using High-Performance Liquid Chromatography (HPLC) according to previous method^[0]^, while volatile apocarotenoids were extracted by Headspace Solid-Phase Microextraction (HS-SPME) and free apocarotenoids were solvent extracted, both aroma compounds were subjected to GC-MS tests^[0]^. Gene expression levels were measured by qRT-PCR using *OfActin* as the reference gene.

For stable transformation in citrus callus, Agrobacterium-mediated transformation was performed on Rm33 callus. The callus line Rm33 was derived from *Citrus reticulata* and maintained in darkness on MT basal medium supplemented with 50 g/L sucrose, 0.5 mg/L 6-benzylaminopurine, and 0.5 mg/L α-naphthaleneacetic acid. Callus pieces (approx. 0.5 g) were co-cultivated with *Agrobacterium* harboring the different *OfCCD4* constructs for 48 hours in the dark. After co-cultivation, tissues were washed and transferred to selection medium containing 50 mg/L hygromycin and 300 mg/L cefotaxime. Transformed calli were subcultured every 3–4 weeks under dark conditions at 25°C. After 2 months of selection, surviving calli were harvested for analysis^[0,0]^. Carotenoid profiles were analyzed via HPLC^[0]^, and volatile aroma compounds were detected using HS-SPME combined with GC-MS ^[0]^. Gene expression levels were measured by qRT-PCR using *CsActin* as the reference gene.

### 4.7. Statistical analysis

All experiments included at least three biological replicates. Data were analyzed using Student’s t-test or one-way ANOVA followed by Tukey’s HSD test in R (v4.3). Graphics were generated with Prism 10 and ggplot2 ^[0]^ in R v.3.6.2.

## Accession Numbers

The raw data for assembly, DNA-seq and RNA-seq have been deposited in the National Genomics Data Center under accession numbers PRJCA055900, PRJCA055898 and PRJCA055907. Detailed information about the sequencing data is listed in Supplementary Table 1, 6 and 8. Genome assembly and annotation of *Osmanthus fragrans* ‘BYYG’ is available at https://figshare.com/s/fb153c4cafffc02127cf.

## Supporting Information

Supporting Information is available from the Wiley Online Library or from the author.

## Acknowledgments

We thank Prof. Deng Xiuxin (Huazhong Agricultural University, Wuhan, China) and Prof. Xu Qiang (Huazhong Agricultural University, Wuhan, China) for providing informative guide and experimental platform for this study. We thank Wuhan Onemore-Tech Co., Ltd. for their assistance with Nanopore ultra-long read sequencing services and the help provided for the T2T genome assembly. We also thank Wuhan Fisha Gene Information Co., Ltd. for their assistance with whole genome resequencing.

## Author Contributions

R.Z. conceived and designed the study. S.L. conducted genome assembly, population genomics, RNA-seq, and genomic variation analyses. S.Y., Z.P., L.Z., L.Z., Y.T., and S.X. performed plant material collection, aroma profiling, carotenoid quantification, and colorimetric (LAB) measurements across cultivars. J.Y. and Q.Y. cloned *OfCCD4* alleles and conducted functional validation assays. X.Z. and W.X. analyzed promoter variation. S.Y. developed the molecular marker. R.Z. supervised the project with support from S.L., Q.X., and X.D. R.Z. and S.L. wrote the manuscript, with contributions from S.Y., S.X., and input from all authors.

## Data Availability Statement

Data are shown in the manuscript or are available from the corresponding author upon request.

