## Supplementary Figure for "T2T Genome and Population Resequencing Reveal *OfCCD4* Alleles Orchestrate Petal Color and Scent in *Osmanthus fragrans*"

**Supplementary Fig. S1 Hi-C interaction map of the *Osmanthus fragrans* genome assembly.**

**Supplementary Fig. S2 BUSCO completeness assessment of the *Osmanthus fragrans* genome.**

**Supplementary Fig. S3 Colorimetric analysis (Lab) and principal component analysis (PCA) of 100 *Osmanthus fragrans* cultivars.**

**Supplementary Fig. S4 Carotenoid and aromatic terpenoid composition in *Osmanthus fragrans* petals.**

**Supplementary Fig. S5 Reads mapping for the 4-bp promoter region of *OfCCD4* in 'Aurantiacus' cultivars.**

**Supplementary Fig. S6 PCR-based detection of *OfCCD4* promoter variants.**

**Supplementary Fig. S7 PCR detection of *OfCCD4* coding sequence (CDS) alleles in non-'Aurantiacus' cultivars.**

**Supplementary Fig. S8 HPLC chromatograms of carotenoids in transgenic *Osmanthus fragrans* petals.**

**Supplementary Fig. S9 GC-MS chromatograms of aroma compounds in transgenic *Osmanthus fragrans* petals.**

**Supplementary Fig. S10 Carotenoid content in transgenic citrus callus.**

**Supplementary Fig. S11 HPLC chromatograms of carotenoids in transgenic *Osmanthus fragrans* petals.**

**Supplementary Fig. S12 GC-MS chromatograms of aroma compounds in transgenic citrus callus.**

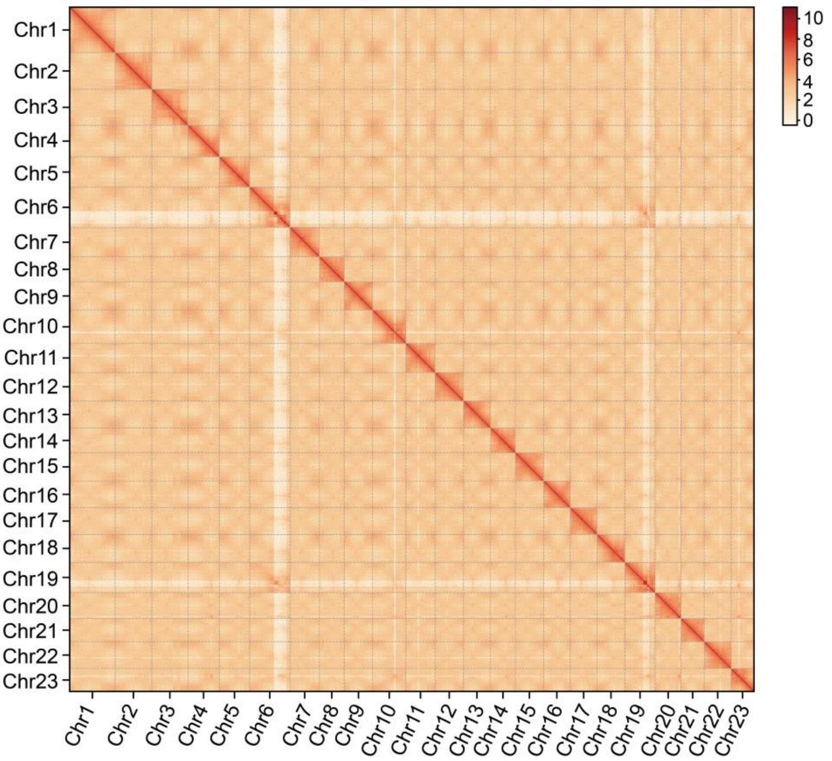


**Supplementary Fig. S1 Hi-C interaction map of the *Osmanthus fragrans* genome assembly.**

The heatmap shows chromatin interaction density, with red indicating higher interaction frequencies.

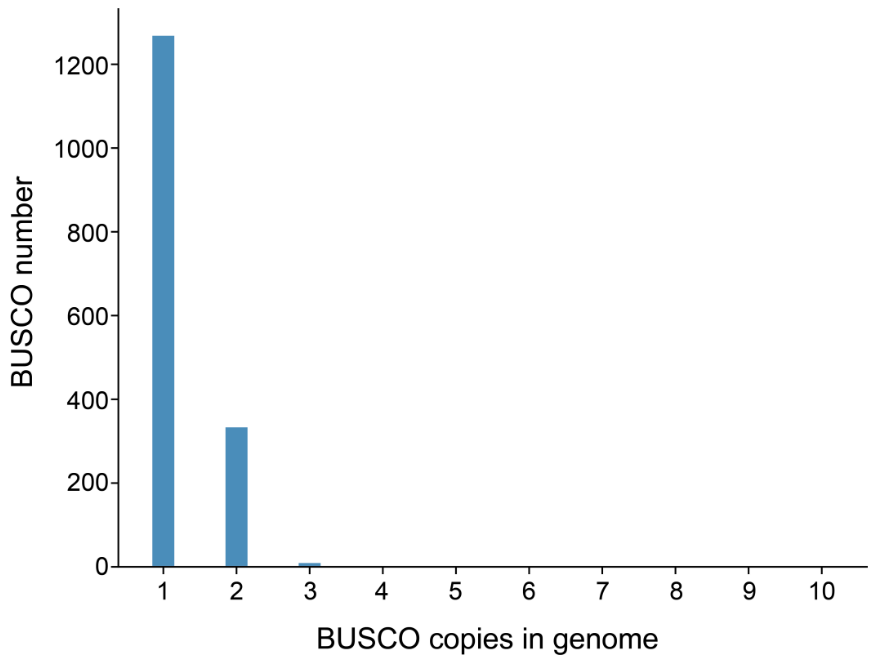


**Supplementary Fig. S2 BUSCO completeness assessment of the *Osmanthus fragrans* genome.**

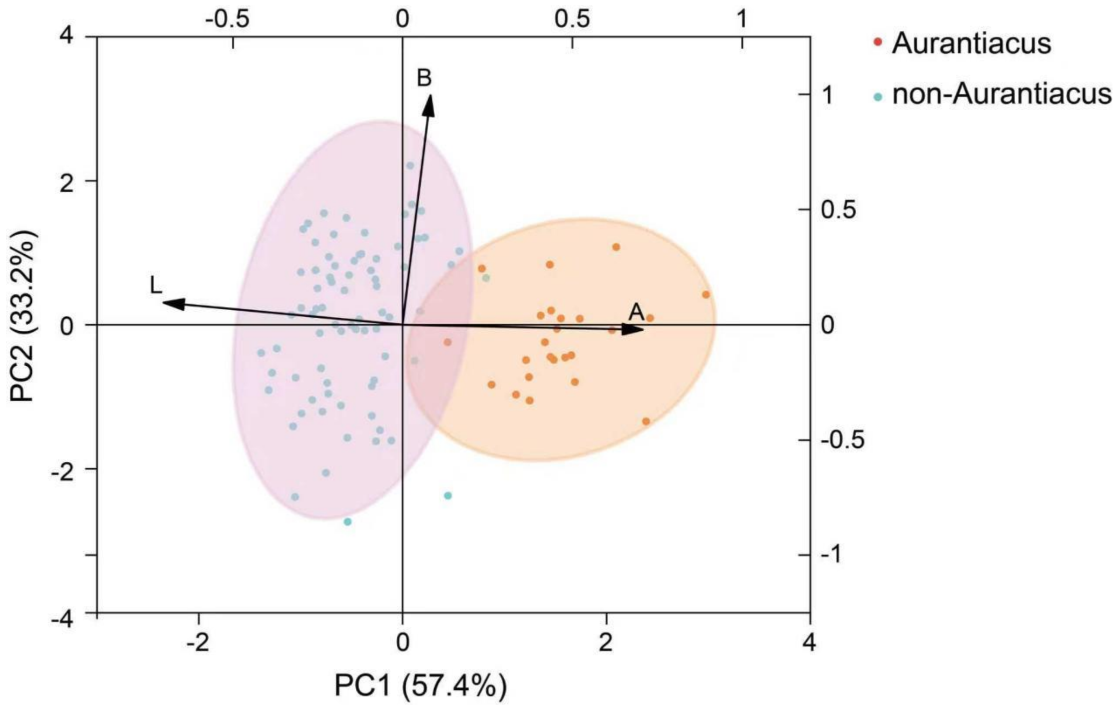


**Supplementary Fig. S3 Colorimetric analysis (Lab) and principal component analysis (PCA) of 100 *Osmanthus fragrans* cultivars.**

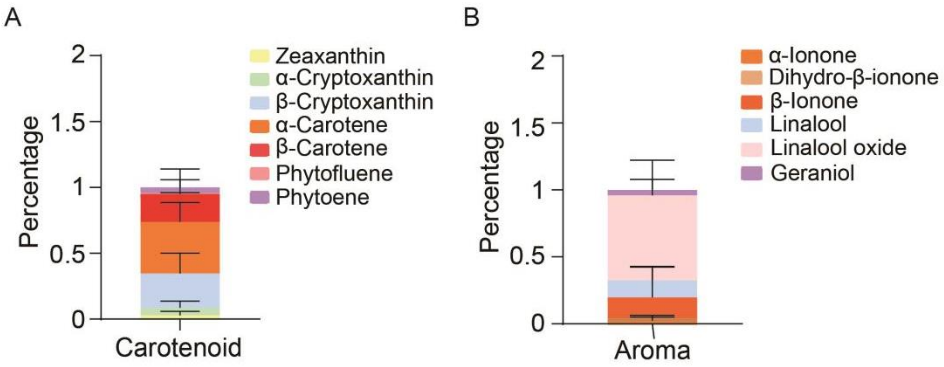


**Supplementary Fig. S4 Carotenoid and aromatic terpenoid composition in *Osmanthus fragrans* petals.**

A, Average proportion of individual carotenoids in petals of 15 'Aurantiacus' cultivars. B, Average proportion of individual aromatic terpenoids in petals of 15 non-'Aurantiacus' cultivars.

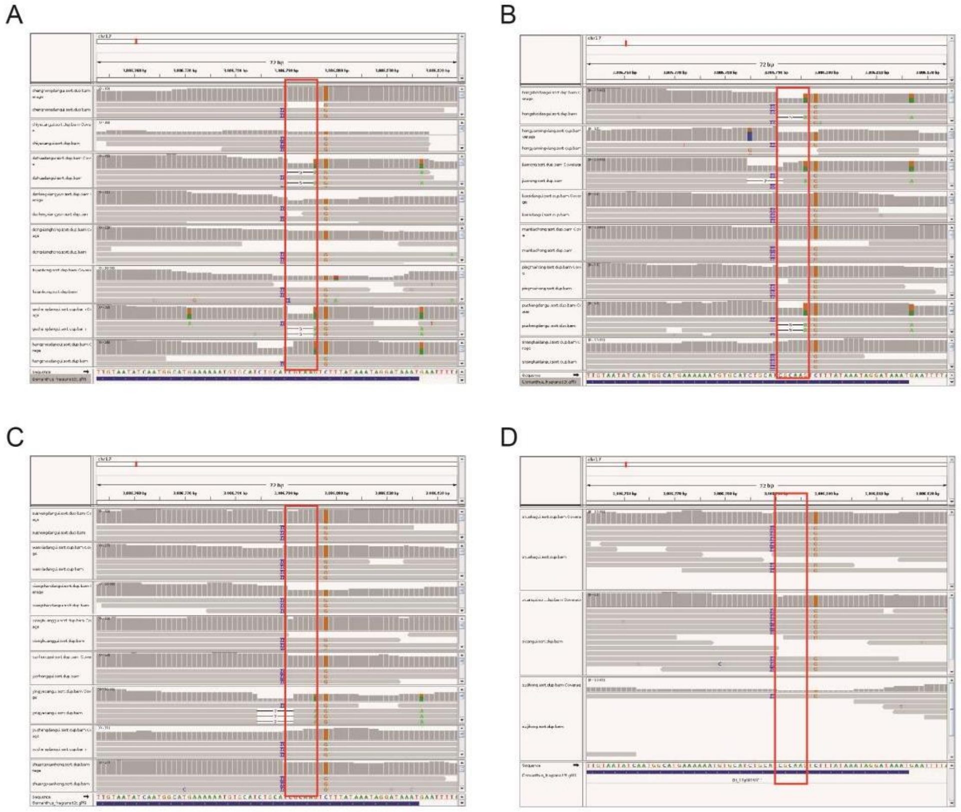


**Supplementary Fig. S5 Reads mapping for the 4-bp promoter region of *OfCCD4* in 'Aurantiacus' cultivars.**

The red box indicates the position of the 4-bp sequence.

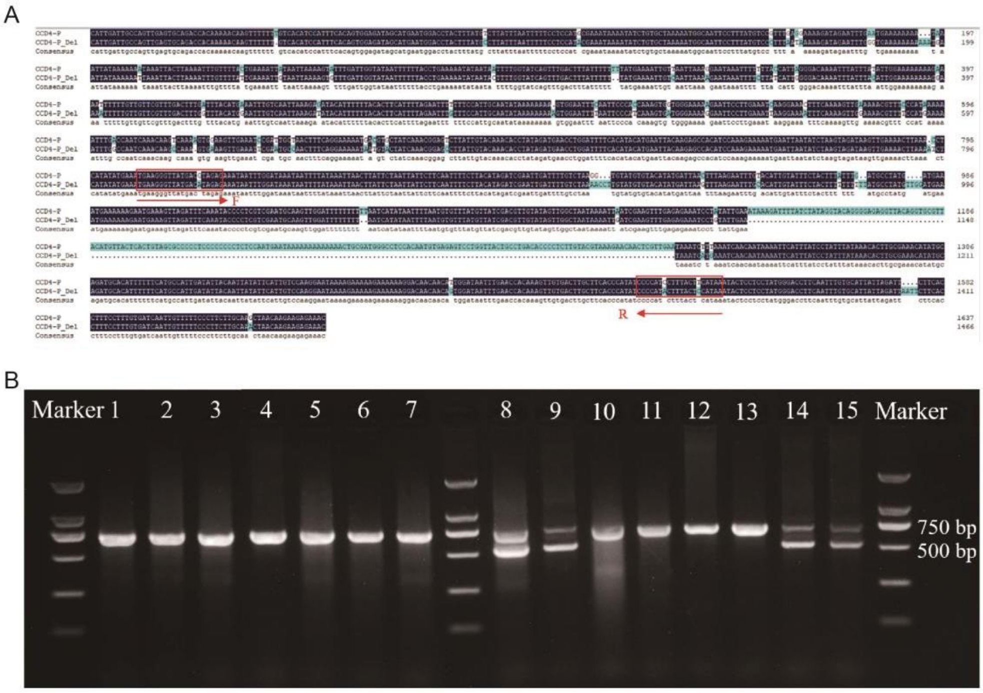


**Supplementary Fig. S6 PCR-based detection of *OfCCD4* promoter variants.**

A, A pair of primers was designed to distinguish between the two promoter types (P and $P^{Del}$). B, The long band represents P, and the short band represents $P^{Del}$. Both variants were detected in 15 ‘Aurantiacus’ cultivars.

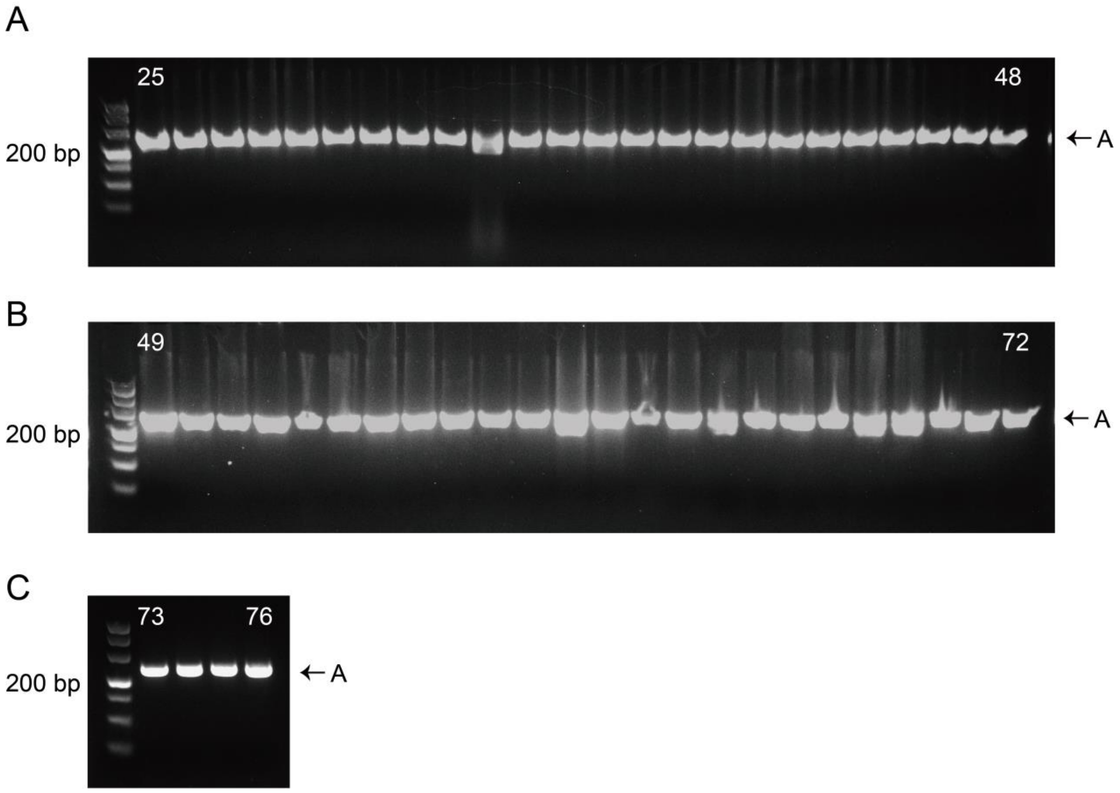


**Supplementary Fig. S7 PCR detection of *OfCCD4* coding sequence (CDS) alleles in non-'Aurantiacus' cultivars.**

Gel electrophoresis shows PCR products for the $a^{Stop}$ and $A/a^{Del}$ alleles. Only the long band (A allele) was detected in non-'Aurantiacus' cultivars.

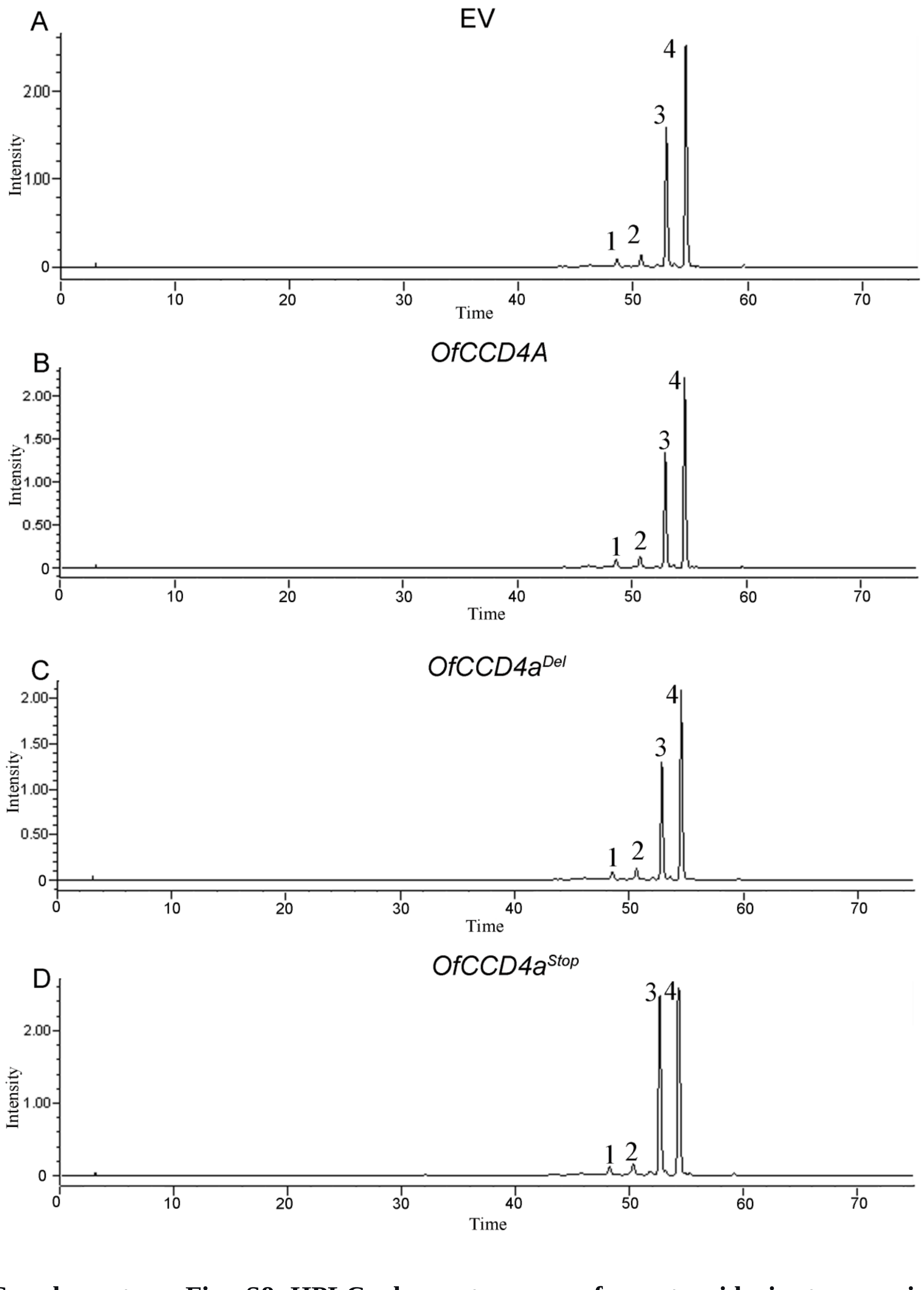


**Supplementary Fig. S8 HPLC chromatograms of carotenoids in transgenic *Osmanthus fragrans* petals.**

Peaks: 1, α-cryptoxanthin; 2, β-cryptoxanthin; 3, α-carotene; 4, β-carotene.

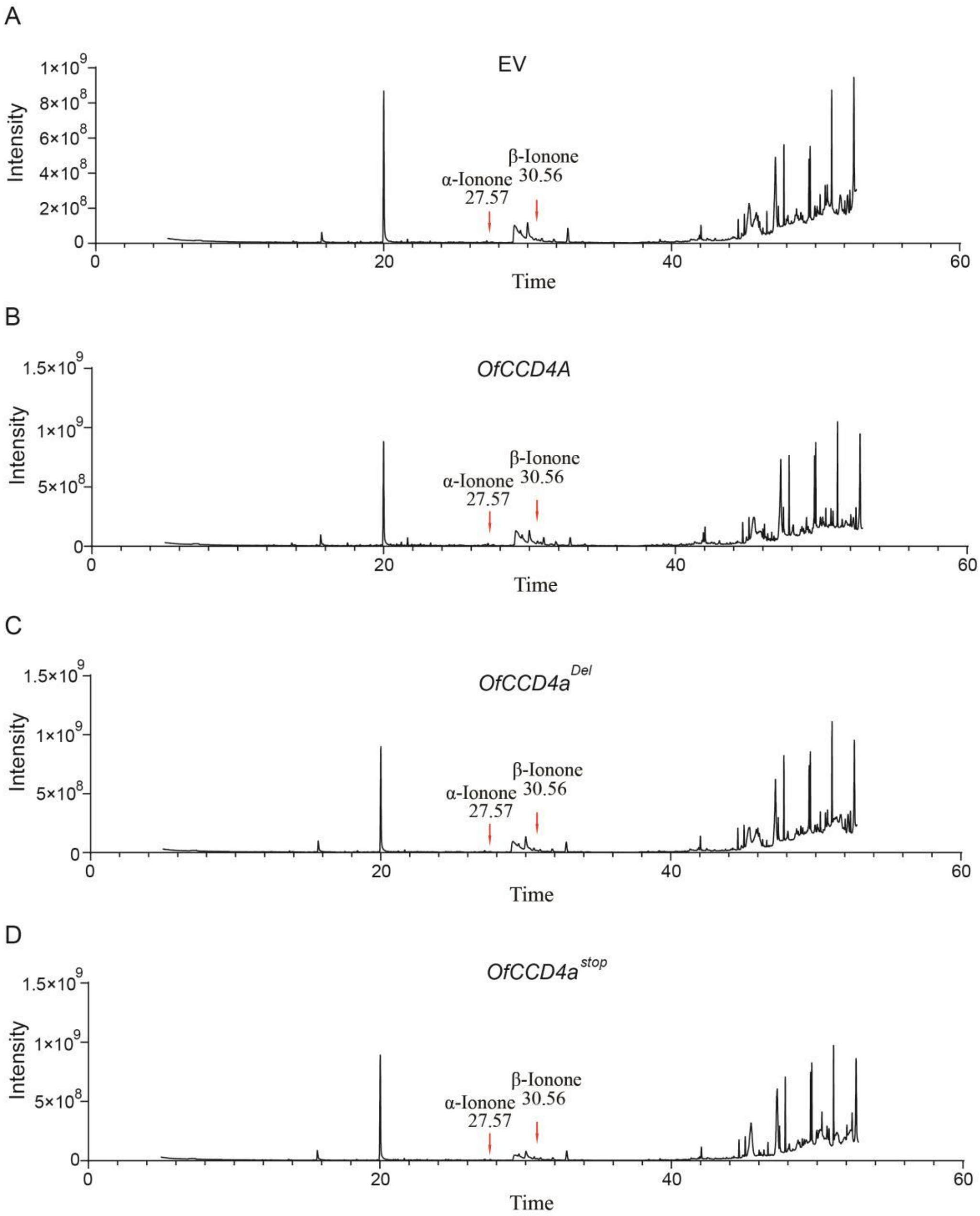


**Supplementary Fig. S9 GC-MS chromatograms of aroma compounds in transgenic *Osmanthus fragrans* petals.**

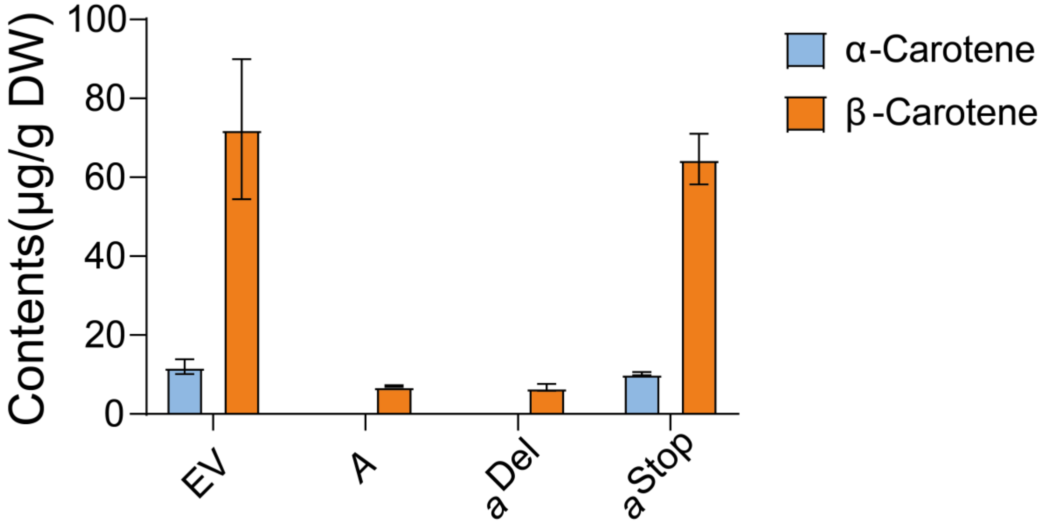


**Supplementary Fig. S10 Carotenoid content in transgenic citrus callus.** Data are presented as mean ± SD of three biological replicates.

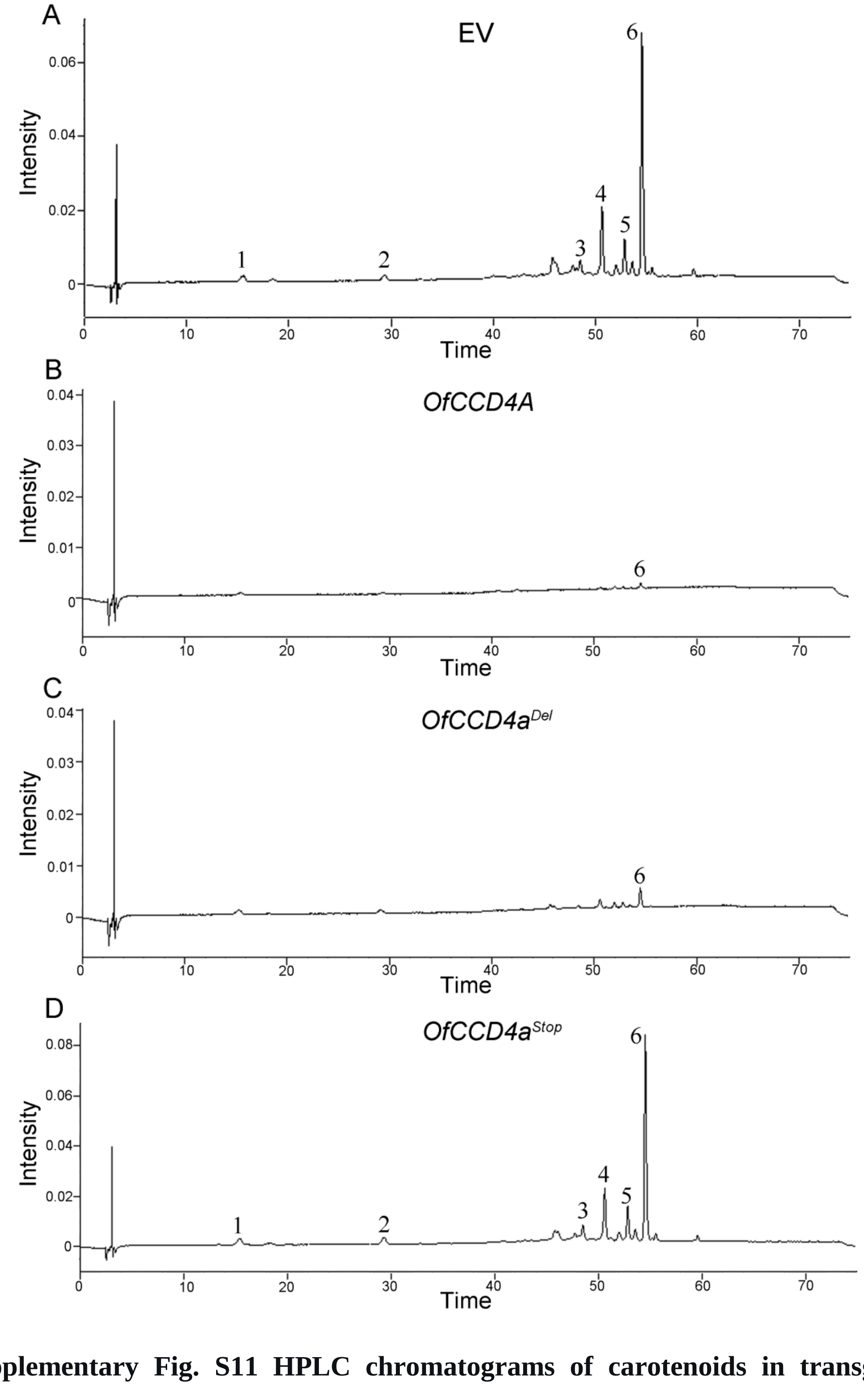


**Supplementary Fig. S11 HPLC chromatograms of carotenoids in transgenic Osmanthus fragrans petals.**

Peaks: 1, violaxanthin; 2, lutein; 3, α-cryptoxanthin; 4, β-cryptoxanthin; 5, α-carotene; 6, β-carotene.

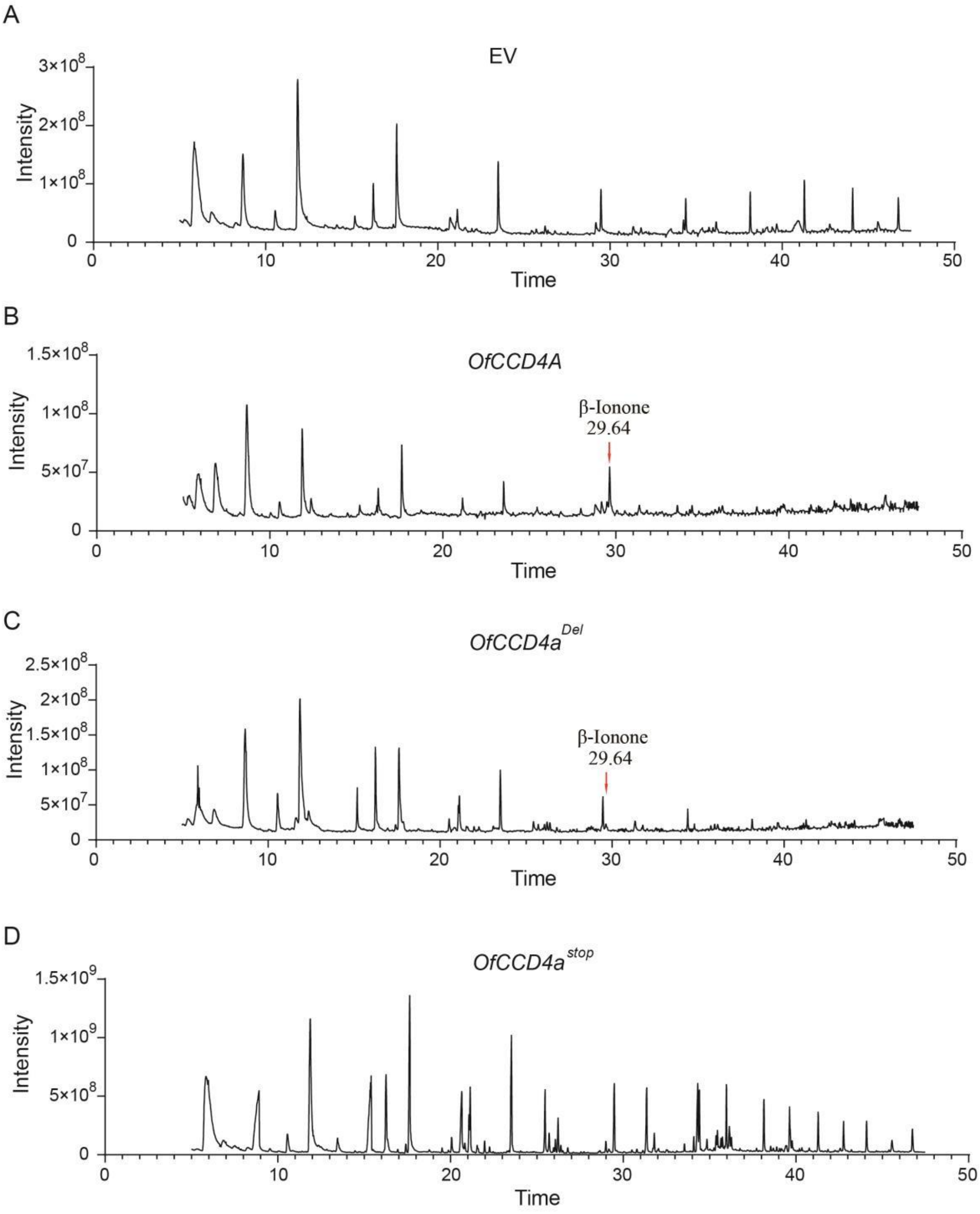


**Supplementary Fig. S12 GC-MS chromatograms of aroma compounds in transgenic citrus callus.**
